# Identification of heat responsive genes in pea stipules and anthers through transcriptional profiling

**DOI:** 10.1101/2021.04.22.440885

**Authors:** Shaoming Huang, Krishna K. Kishore, Reddy V.B. Lachagari, Navajeet Chakravartty, Rosalind A. Bueckert, Bunyamin Tar’an, Thomas D. Warkentin

## Abstract

Field pea (*Pisum sativum* L.), a cool-season legume crop, is known for poor heat tolerance. Our previous work identified PR11-2 and PR11-90 as heat tolerant and susceptible lines in a recombinant inbred population. CDC Amarillo, a Canadian elite pea variety, was considered as another heat tolerant variety based on its similar field performance as PR11-2. This study aimed to characterize the differential transcription. Plants of these three varieties were stressed for 3h at 38°C prior to self-pollination, and RNAs from heat stressed anthers and stipules on the same flowering node were extracted and sequenced via the Illumina NovaSeq platform for the characterization of heat responsive genes. *In silico* results were further validated by qPCR assay. Differentially expressed genes (DEGs) were identified at log2 fold change, the three varieties shared 588 DEGs which were up-regulated and 220 genes which were down-regulated in anthers when subjected to heat treatment. In stipules, 879 DEGs (463/416 upregulation/downregulation) were consistent among varieties. The above heat-induced genes of the two plant organs were related to several biological processes i.e., response to heat, protein folding and DNA templated transcription. Ten gene ontology (GO) terms were over-represented in the consistently down-regulated DEGs of the two organs, and these terms were mainly related to cell wall macromolecule metabolism, lipid transport, lipid localization, and lipid metabolic processes. GO enrichment analysis on distinct DEGs of individual pea varieties suggested that heat affected biological processes were dynamic, and variety distinct responses provide insight into molecular mechanisms of heat-tolerance response. Several biological processes, e.g., cellular response to DNA damage stimulus in stipule, electron transport chain in anther that were only observed in heat induced PR11-2 and CDC Amarillo, and their relevance to field pea heat tolerance is worth further validation.

## Introduction

Human activities have contributed approximately 1°C temperature increase globally since the Industrial Age, and are predicted to cause another 0.5-1°C increase in the period between 2030 and 2052 according to current greenhouse gas emission rates [1]. The evidence of the rising temperature causing lowered grain production was reported in the three major crops, maize, wheat, and rice [2]. Heat stress (HS) also limits the production on legume crops including pea. In Canada, where its pea production accounts for one third of the global production, lowered grain yield was observed in summers when the maximum temperature exceeded 28 °C during flowering, or the seasonal temperature was over 17.5 °C [3, 4]. Because of the concern about a warming summer in North America, physiological studies on HS related damage on field pea, particularly the reproductive plant parts, have been conducted in the last decade. When pea plants at anthesis were exposed to 36/18°C day/night for 7 days in a growth chamber, the pollen germination percentage, pollen tube length, pod length, seed number per pod, and the seed–ovule ratio dropped dramatically compared to pea exposed to normal conditions of 24/18°C [5]. In addition, HS reduced both pollen and ovule viability, but pollen appeared to be more heat susceptible [6]. In terms of pea breeding, progress has also been made in the characterization of heat tolerance based on field trials. A longer duration from sowing to flowering termination, and greater pod production per plant contributed to increased grain yield potential at both hot and normal conditions, and several stable quantitative trait loci were characterized related to flowering and yield component traits [4]. Lodging resistance and the semi-leafless leaf type resulted in a cooler pea canopy and greater yield potential [7]. Additionally, the authors further characterized putative genomic loci of heat responsive traits, e.g., canopy temperature, pod number and chlorophyll concentration, via a pea genome wide association mapping study [8].

The discovery of heat responsive genes started with the characterization of heat shock protein (HSP) genes and their transcription factors (HSFs). Findings in this aspect were firstly well documented in *Arabidopsis thaliana*. In addition to the 21 known HSFs [9], the Arabidopsis heat response is partly mediated by 13 HSP20s [10], 18 HSP70s [11], seven HSP90s [12], and up to eight members of the HSP100s [13]. The gene family of HSP20 was most highly expressed under HS, followed by the gene family of HSP70 and HSP90, and the gene family of HSP100 was not responsive to heat stress [14]. Subsequent studies on the global transcriptome profiling under HS revealed that heat responsive genes could expand to those other genes involved in plant hormone biosynthesis and signaling, calcium and sugar signaling, primary and secondary metabolism [15–17]. Cell wall and secondary metabolite pathways were also highly affected under HS in lentil [18]. However, both the number of up- and down-regulated genes and the ratio of up- and down-regulated genes under HS varied among the above mentioned studies depending on HS treatments, plant species, genotypes and different plant organs used for RNA isolation.

Research on heat responsive gene discovery in pea is limited to the findings of HSP genes. Among the reported pea HSP genes, the expression of *PsHSP 18.1* and *PsHSP71.2* genes appeared to be heat inducible [19, 20]. The relation of HSPs to heat tolerance was subsequently confirmed as the induction of these HSP genes improved survival rate of pea seedlings and mature plants at high temperature [21]. Moreover, several HSP genes had greater heat-induced expression in one of the heat tolerant cultivars, Acc.623, than in one susceptible variety Acc.476. The conclusion that the heat tolerant variety, in general, outperformed the susceptible variety in terms of HSP heat induction threshold is still in question because this study did not conduct a full comparison of HSP heat induced expression patterns among the pair of pea heat tolerant and susceptible varieties.

Although lacking the reference genome previously, transcriptome profiling via RNA-seq studies were carried out in pea over the last decade, mainly focusing on the mining of genetic markers. The first pea transcriptome reference was developed using next generation sequencing with the Roche/454 platform [22]. Later Illumina high-throughput sequencing was applied to sequence 23 cDNA libraries from multiple tissues of the Australian field pea cultivars Kaspa and Parafield [23]. A large proportion of the assembled contigs were expressed in both cultivars. To date, no transcriptome-wide mapping of pea response to HS has been conducted, but this method was utilized in the discovery of responsive genes in field pea seed aging [24], root nodulation [25] and most recently in water-logging stress studies [26, 27]. The utilization of RNA-seq technique in pea HS research allows for the genome-wide mining of heat responsive genes and the global description of the complex regulatory pathway in the protection against HS at the cellular level, as well as comparative analysis of genes responsive to HS among different pea varieties, or between pea and other crop species. Thus, the objectives of this research included 1) characterization of additional gene response toward high temperature besides previously characterized pea HSP genes; 2) comparative analysis of heat responsive gene expression differences between anthers and stipules, as well as between heat tolerant and heat susceptible varieties.

## Methods

### Plant materials

Three pea varieties were used as plant material for this experiment, that is, PR11-2 (heat tolerant variety), PR11-90 (heat susceptible variety) and CDC Amarillo (check variety). PR11-2 and PR11-90 are recombinant inbred lines from the population PR11, which was derived from the cross CDC Centennial/CDC Sage made in 2008 at the Crop Development Centre (CDC), University of Saskatchewan [4]. CDC Centennial was developed at CDC. It is a high yielding yellow pea cultivar with semi-leafless leaf type with moderately large seeds [28]. CDC Sage is a high yielding cultivar from the CDC with green cotyledons and medium-small seeds [29]. PR11-2 and PR11-90 have white flowers and green cotyledons, but PR11-2 has greater pod number per plant, longer flowering duration and greater grain yield than PR11-90 based on field trials at both normal and hot conditions, thus PR11-2 is considered to have better heat tolerance than PR11-90. CDC Amarillo [30], a yellow pea variety and one of the best yielding varieties in western Canada, was included as a check. Because CDC Amarillo has similar field performance as PR11-2 in our field test at normal and heat stressful conditions (Table 1), it is also considered as heat tolerant compared with PR11-90.

**Table 1.** Characteristics of flowering and yield-related traits of PR11-2, PR11-90 and CDC Amarillo at normal and late seeding trials in 2017-2019 at Saskatoon, Canada.

| variety | seeding date | DTF | DOF | RNN | PN | SNPP | TSW(g) | plot yield(kg/ha) |
| --- | --- | --- | --- | --- | --- | --- | --- | --- |
| PR11-2 | normal | 56.9 | 15.6 | 5.4 | 7.7 | 4.1 | 215.4 | 2827.7 |
|  | late | 51.9 | 12.7 | 4.8 | 7.4 | 3.5 | 216.6 | 2665.2 |
| PR11-90 | normal | 48.0 | 17.7 | 4.5 | 7.1 | 5.8 | 200.9 | 2288.9 |
|  | late | 47.7 | 13.0 | 3.3 | 5.2 | 4.8 | 177.6 | 1144.9 |
| CDC Amarillo | normal | 56.5 | 14.5 | 5.0 | 7.6 | 4.6 | 236.4 | 3064.5 |
|  | late | 51.4 | 13.6 | 4.4 | 7.1 | 3.9 | 200.9 | 2734.2 |

Note: DTF, days to flowering; DOF, duration of flowering; RNN, reproductive node number on main-stem; PN, pod number on main-stem; SNPP, seed number per pod; TSW, thousand seed weight (g). Late seeding trial is more heat stressful trial.

### Experimental design

A randomized complete block design experiment utilizing the three varieties with three biological reps and two temperature treatments was carried out in a phytotron chamber in the Agriculture Building, University of Saskatchewan. Temperature treatments consisted of control temperature treatment 24/16°C, 16/8h day/night, and high temperature treatment 38/16°C, 16/8h day/night. Three seeds of each variety were planted in individual 3.8 L pots containing Sunshine mix #4 (Sun Gro, Seba Beach, AB, Canada). The three plants in one pot were bulked later as one biological replication. Starting from 1 week after crop emergence, the plants were watered every 2-3 days based on the growth stage and water use. Once a week, a quick release fertilizer (20 N:20 P_2_O_5_:20 K_2_O) prepared at a concentration of 3 g L^-1^ was applied at a rate of 100 ml per pot starting 1 week after emergence. At the stage when plants developed the first flower bud, but before anther dehiscence, pots of all varieties in the heat treatment group were transferred to a 38°C chamber for 3h. Then all the anthers and stipules on the first flowering node of the three plants within one pot were sampled from both normal and high temperature treatments, and then were freshly frozen in liquid nitrogen and kept at −80°C for storage.

### RNA extraction and RNA integrity check

The whole experiment constituted a library of 36 samples from three varieties, two plant organs, two temperature treatments and three biological reps, as detailed in the previous section. For each organ sample, the extraction of total RNA was conducted using QIAGEN RNeasy plant mini kit from QIAGEN Inc, and then a further clean-up step by digesting any remaining DNA contaminant was carried out using QIAGEN RNase-free DNase set. The quantity of extracted RNA sample was then determined by evaluating optical density at 260 nm and the OD260/OD280 absorption ratio using NanoDrop 8000 UV spectrophotometer. The integrity of all 36 RNA samples were profiled for integrity via Bioanalyzer 2100 according to the manufacturer’s manual, and all RNA samples had integrity scores in the range of 9-10 on the scale of 0-10, which passed the integrity standard for sequencing.

### RNA-Seq protocol

Construction of cDNA libraries and subsequent sequencing was done at MedGenome Inc (https://www.medgenome.com, Foster City, CA, USA).

### Raw data processing and sequencing read alignment

In the pre-processing step of the raw reads, the adapter sequences and low-quality bases were trimmed using AdpaterRemoval-V2 [31]. From the preprocessed reads, ribosomal RNA sequences were removed by aligning the reads with SILVA database [32] using Bowtie2_v2.2.9 [33]. The remaining reads were aligned to the pea reference genome (Pisum_sativum_v1a.fa) and gene model (Pisum_sativum_v1a_genes.gff3) [34]. The alignment was preformed using STAR_v2.5.3a [35].

### Differential gene expression analysis and annotation

Firstly, a homology search was executed for all 44,756 gene sequences against UniProt plant [36] using Diamond_v0.9.3.104 [37]. Out of 44,756 genes, 33,669 genes were annotated based on tophit. Then for each variety, the differential expression analysis between heat-treatment (3H 38°C) and control (22°C) was conducted via cuffdiff program in cufflinks package_v 2.2.1 [38]. Log2 fold change (FC) cutoff 2/-2 and p-value cutoffs 0.01 were used separately as cutoffs for up and down regulated genes to characterize differentially expressed genes (DEGs). The unit of measurement used by Cufflinks to estimate transcript abundance is fragments per kilobase of transcript per million mapped reads.

### Quantitative real-time PCR validation

To validate the correctness of above DEG results analysed *in silico*, qPCR bench assay was conducted to test the result consistency between the two methods. Eleven random genes were originally selected from the pea genome and primers were designed for each gene via IDT Primer quest tool (Integrated DNA Technologies Inc) according to the following criteria, i.e., Tm of 62 ± 1°C, PCR amplicon lengths of 90-120 bp, primer length of 20-22 bp, and GC content of 45-55%. A series of 10-time cDNA dilutions on PR11-90_control leaf cDNA library was made for primer efficiency test. And primer efficiency (%) of each gene was equaled to (10^―1/*slope*^ ―1) ∗ 100, and all primers had their efficiency rates between 90-110% and qualified for assay use (S1 Fig).

Subsequently, the relative expression of the 11 genes was separately quantified among 18 stipule samples and the 18 anther samples, which were used for RNA sequencing. RT-qPCR data were analyzed according to the comparative 2^-ΔΔCt^ method [39], where ΔCt = (Ct of gene of interest – Ct of reference gene). The relative gene expression change was compared between qPCR bench assay and RNA-Seq on stipules and anthers separately.

### Gene ontology (GO) enrichment analysis on DEGs

Further comparative analyses on DEGs were conducted among the three pea varieties with different heat tolerances, and between anther (reproductive plant organ) and stipule (vegetative plant organ). The results were output in Venn diagram via online software (http://bioinformatics.psb.ugent.be/webtools/Venn/). Subsequently, GO terms of heat responsive genes were tested against the pea reference transcriptome (Pisum_sativum_v1a_GO, database was retrieved in November, 2020) via agriGO v2.0 [40], and significant GO terms in biological processes were filtered using hypergeometric test method at FDR adjusted p value<0.01.

## Results

### Sequencing quality assessment

To understand transcriptional reprogramming of field pea in response to heat stress, we performed deep RNA sequencing of stipule and anther organs subjected to 38°C for 3h among three varieties using the NovoSeq sequencing platform. The sequencing platform produced a high confidence sequencing output with a <2% maximum read error rate among the 36 libraries. After removing the error reads, the anther libraries had an average of 84 million 100 bp paired-end reads across the three varieties (Table 2). Stipule libraries resulted in a similar average of 88 million reads (Table 3). Both types of libraries outputted high sequencing depths for the global transcriptome analysis as compared with previous pea transcriptome studies [23, 25]. Subsequently the reads were mapped to the pea reference genome [34], and nearly all reads were successfully aligned to the pea genome, which also implies good quality of the deep sequencing.

**Table 2.**
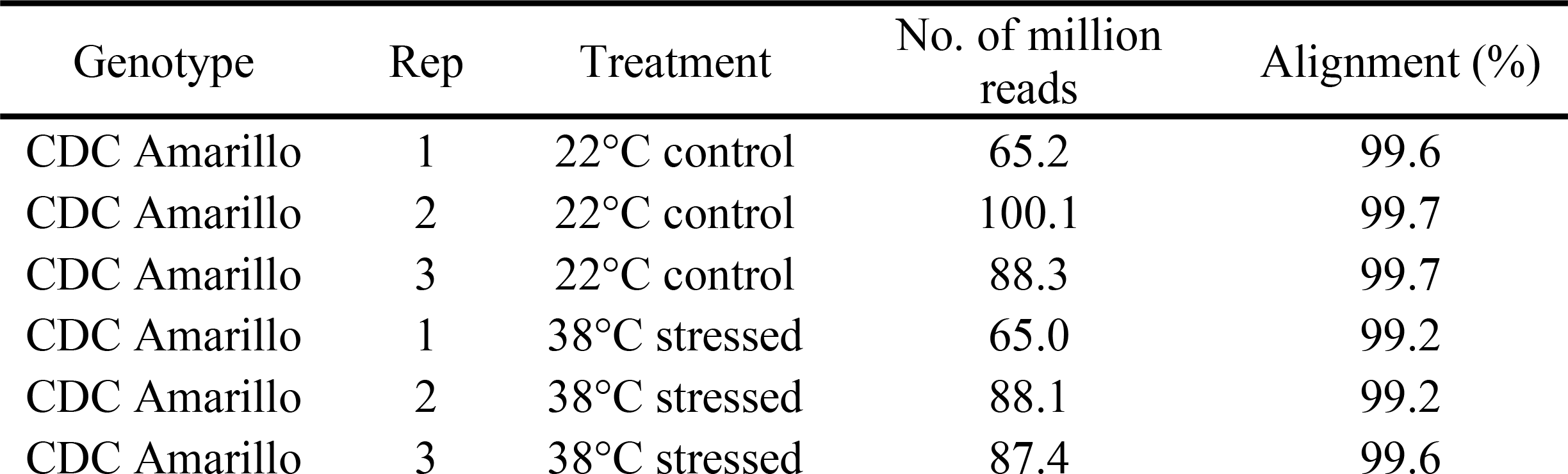
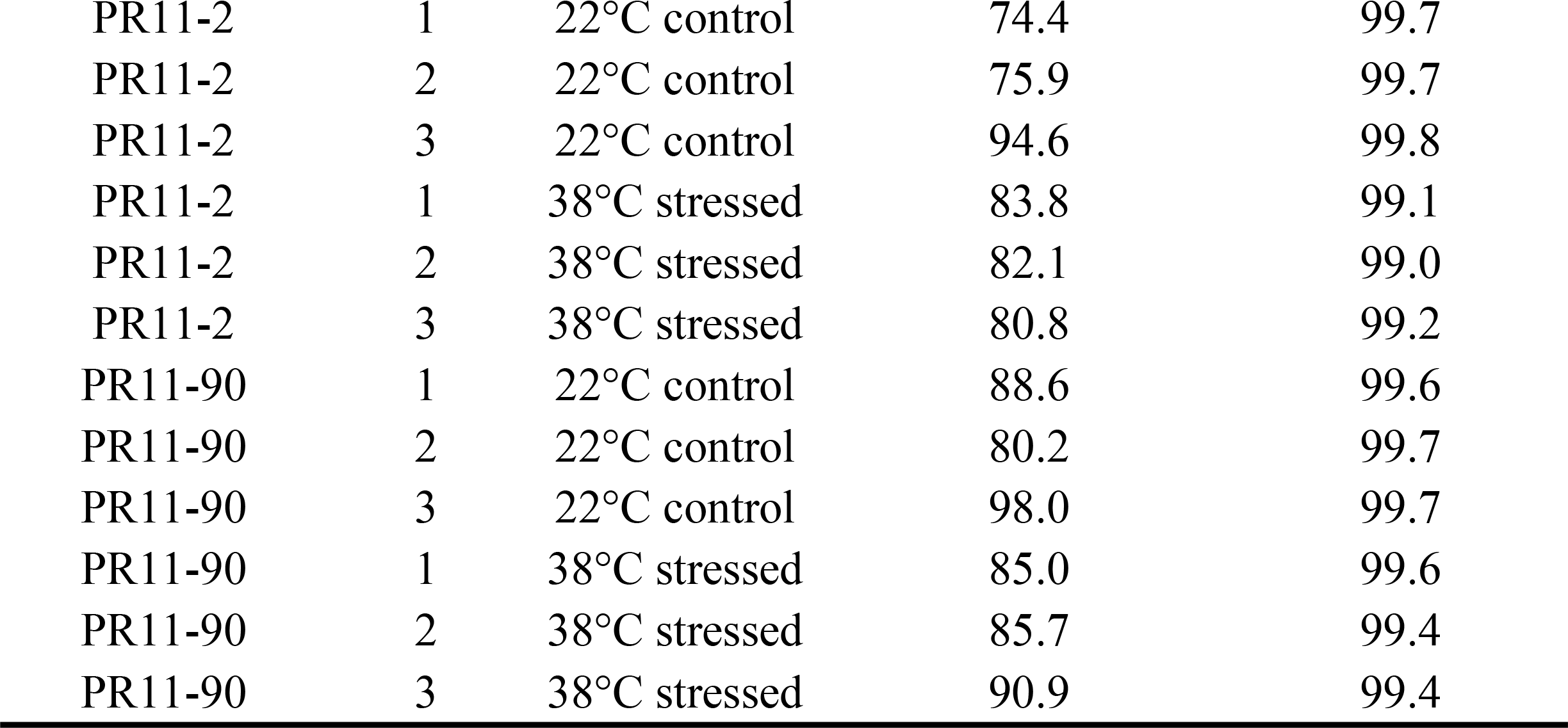
Summary of sequencing depth and percentage of sequencing reads aligning to pea genome on anther samples.

| Genotype | Rep | Treatment | No. of million reads | Alignment (%) |
| --- | --- | --- | --- | --- |
| CDC Amarillo | 1 | 22°C control | 65.2 | 99.6 |
| CDC Amarillo | 2 | 22°C control | 100.1 | 99.7 |
| CDC Amarillo | 3 | 22°C control | 88.3 | 99.7 |
| CDC Amarillo | 1 | 38°C stressed | 65.0 | 99.2 |
| CDC Amarillo | 2 | 38°C stressed | 88.1 | 99.2 |
| CDC Amarillo | 3 | 38°C stressed | 87.4 | 99.6 |
| PR11-2 | 1 | 22°C control | 74.4 | 99.7 |
| PR11-2 | 2 | 22°C control | 75.9 | 99.7 |
| PR11-2 | 3 | 22°C control | 94.6 | 99.8 |
| PR11-2 | 1 | 38°C stressed | 83.8 | 99.1 |
| PR11-2 | 2 | 38°C stressed | 82.1 | 99.0 |
| PR11-2 | 3 | 38°C stressed | 80.8 | 99.2 |
| PR11-90 | 1 | 22°C control | 88.6 | 99.6 |
| PR11-90 | 2 | 22°C control | 80.2 | 99.7 |
| PR11-90 | 3 | 22°C control | 98.0 | 99.7 |
| PR11-90 | 1 | 38°C stressed | 85.0 | 99.6 |
| PR11-90 | 2 | 38°C stressed | 85.7 | 99.4 |
| PR11-90 | 3 | 38°C stressed | 90.9 | 99.4 |

**Table 3.** Summary of sequencing depth and percentage of sequencing reads aligning with pea genome on stipule samples.

| Genotype | Rep | Treatment | No. of million reads | Alignment (%) |
| --- | --- | --- | --- | --- |
| CDC Amarillo | 1 | 22°C control | 101.7 | 99.3 |
| CDC Amarillo | 2 | 22°C control | 92.1 | 99.5 |
| CDC Amarillo | 3 | 22°C control | 100.1 | 99.5 |
| CDC Amarillo | 1 | 38°C stressed | 82.6 | 97.3 |
| CDC Amarillo | 2 | 38°C stressed | 78.3 | 98.4 |
| CDC Amarillo | 3 | 38°C stressed | 80.4 | 99.1 |
| PR11-2 | 1 | 22°C control | 90.3 | 99.6 |
| PR11-2 | 2 | 22°C control | 90.4 | 99.3 |
| PR11-2 | 3 | 22°C control | 87.9 | 99.6 |
| PR11-2 | 1 | 38°C stressed | 85.2 | 98.8 |
| PR11-2 | 2 | 38°C stressed | 84.7 | 99.3 |
| PR11-2 | 3 | 38°C stressed | 106.4 | 99.3 |
| PR11-90 | 1 | 22°C control | 77.5 | 99.0 |
| PR11-90 | 2 | 22°C control | 109.8 | 99.6 |
| PR11-90 | 3 | 22°C control | 80.4 | 99.5 |
| PR11-90 | 1 | 38°C stressed | 74.7 | 98.9 |
| PR11-90 | 2 | 38°C stressed | 66.5 | 99.0 |
| PR11-90 | 3 | 38°C stressed | 107.7 | 99.3 |

### DEG analysis validation

The stipule expression response (log2 FC) of the 11 randomly selected genes between heat treatment and control temperature were characterized via cuffdiff program and qPCR respectively, and the results are shown in a heat map. Nine genes out of the eleven, had consistent heat responses between qPCR and RNA-Seq result *in silico* (Fig 1), implying a good quality of RNA-Seq analysis. And the significant correlation (R^2^ = 0.97) between bench result and *in silico* result further confirmed a correct analysis via cuffdiff program (Fig 2a). Two genes had some unmatched results between the two methods. 0s3930g0040 displayed a consistently up-regulated expression via qPCR among the three pea varieties when subjected to heat treatment (Fig 1), whereas for the *in silico* result only CDC Amarillo had the same trend. From *in silico* result, 5g006560 demonstrated a consistent downregulation towards HS in all varieties; whereas in qPCR result, only PR11-90 had the similar trend. Likewise, among anther samples, a high consistency was found between bench results and cuffdiff result (Fig 1 and 2b). Unmatched results were mainly observed on gene 5g006560. The significantly high correlation (R^2^ = 0.93) between the two methods confirmed the correctness of the analyses. The qPCR results successfully validated the correctness of RNA-Seq analysis on both anther and stipule samples.

**Fig 1.**
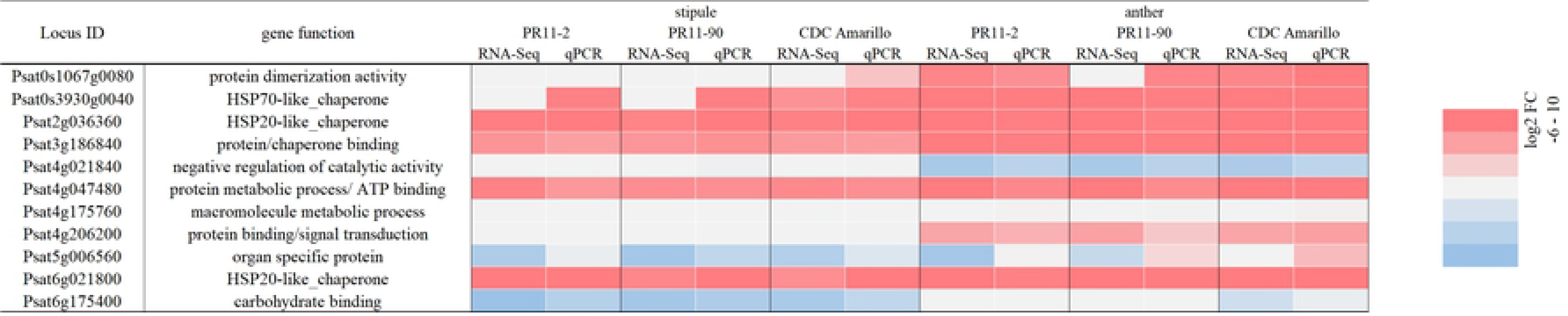
Transcriptional heat response heatmap of 11 randomly selected genes in the pea genome via qPCR and cuffdiff *in silico* methods. FC values are the average log2 (FC) across three biological reps; red color is for upregulation and blue color is for downregulation.

**Fig 2.**
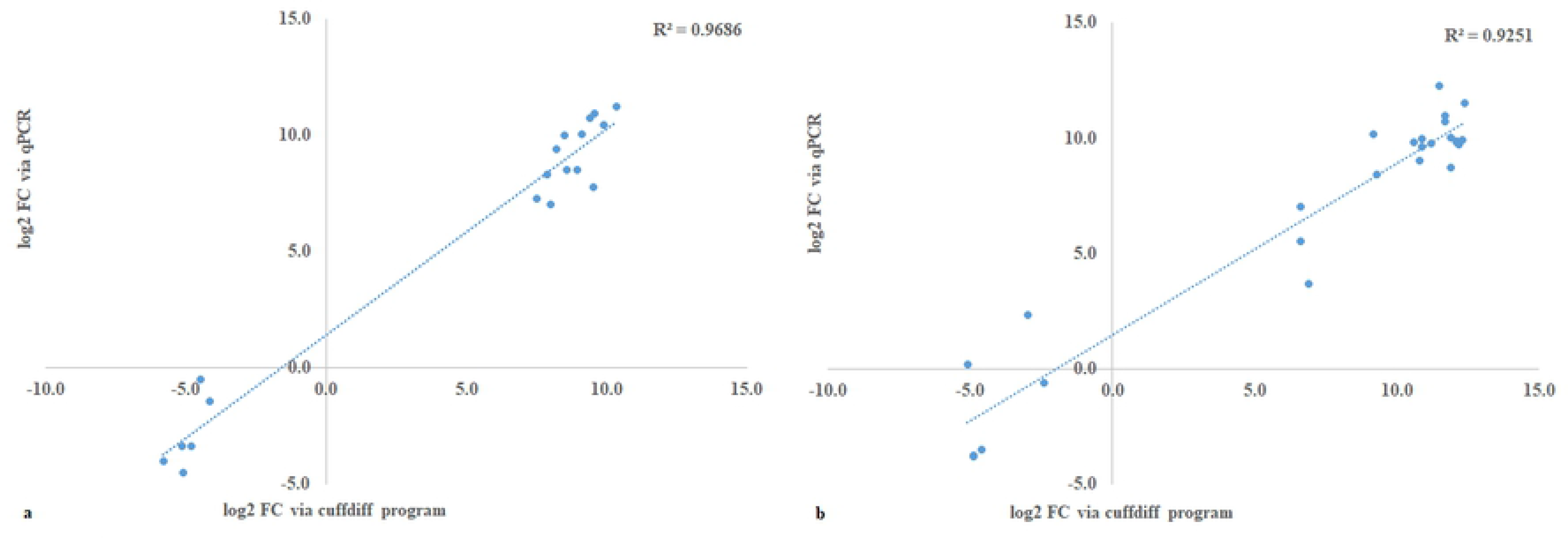
Gene expression result correlation on stipule samples (panel a) and anther samples (panel b) between qPCR and cuffdiff program.

### Global comparisons of HS related transcriptomes between stipules and anthers among three pea varieties

To gain the knowledge on gene response to heat treatment, genes whose expression differed between HS and control temperature at log2ǀFCǀ≥2 were characterized as heat responsive genes. A total of 3565 responsive genes were identified in anthers, among which 2322 genes had greater expression and 1243 had lower expression in heat treatment compared to control temperature. Stipules on the same flowering node had 4381 responsive genes, with 1886 up-regulated genes and 2495 down-regulated genes. Among anther transcriptomes of the three varieties, the number of genes that were up-regulated under HS was almost twice the number of down-regulated genes (Fig 3). The three varieties shared 588 genes with up-regulated expression under HS, which comprised of 25% up-regulated genes in total. The overlap between PR11-2 and PR11-90, where the two varieties were derived from the same recombinant inbred population, accounted for a higher proportion (∼70% in PR11-2 and ∼60% in PR11-90). CDC Amarillo, which has a different genetic background, contributed a major group of DEGs that were distinctly up-regulated. Among the 1343 genes whose expression was inhibited, 220 genes were found common among the three varieties.

**Fig 3.**
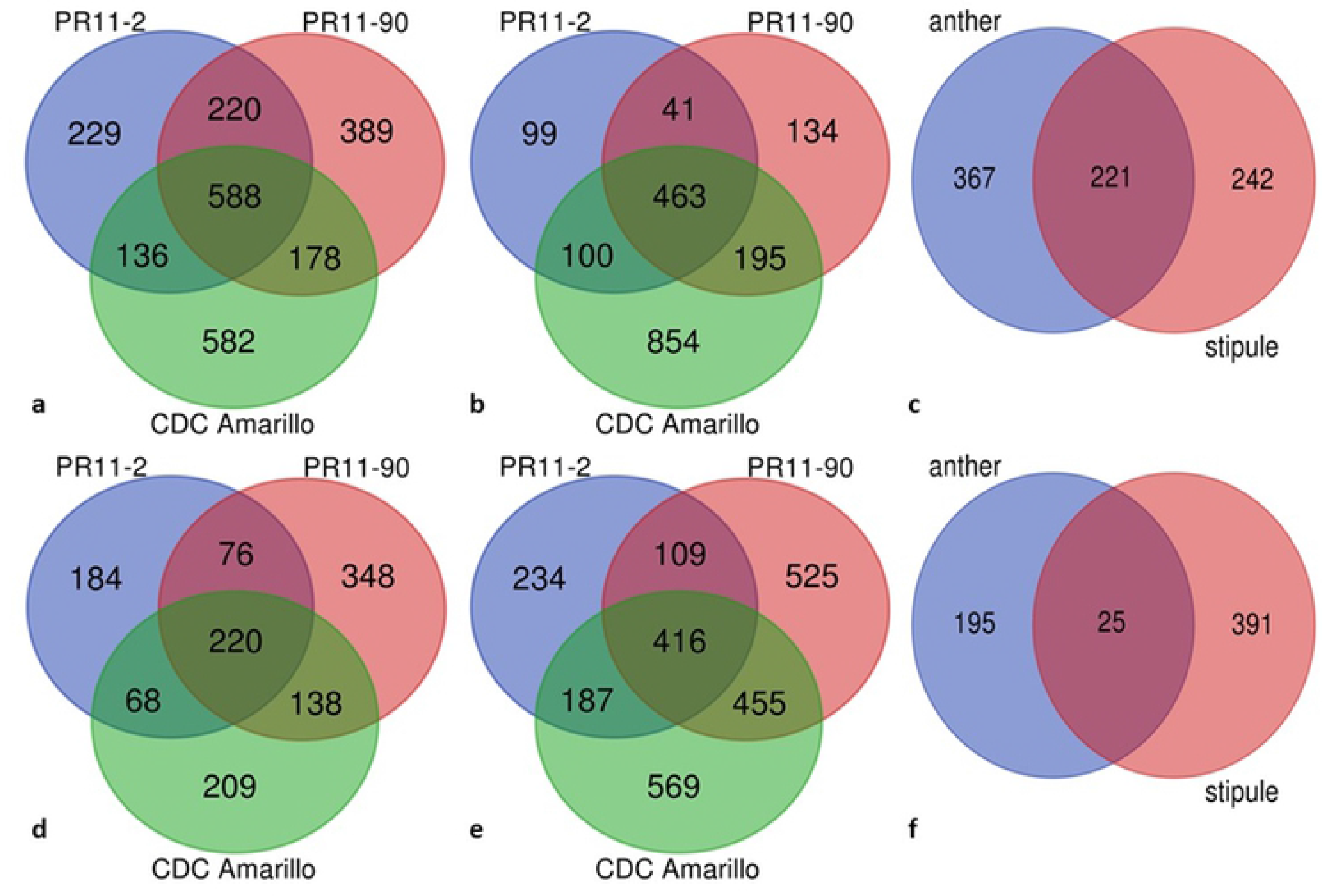
Venn diagram showing the number of common and specific differentially expressed genes (log2 ǀFCǀ ≥ 2; false discovery rate < 0.05) at 3h 38°C heat treatment among three pea varieties, and between anther and stipule on the same node. Panel a-c are for up-regulated genes (from the left to right are anther, stipule and comparison between the two. Panel d-f are for down-regulated genes in the same order mentioned above.

Whereas among the surrounding stipule leaf transcriptomes, the pattern was opposite compared to the anther transcriptome, i.e., a greater number of genes were down-regulated in the heat treatment. The result revealed a different heat response in stipules compared to anthers. Still, there were common DEGs between anthers and stipules, 220 common DEGs with their up-regulated expression and 25 DEGs with down-regulated expression. Respective gene ontology (GO) enrichment analysis of the two groups of DEGs was conducted to cluster their functions in plant biological processes, and results were elucidated in the section below on GO analysis.

Among the three varieties, PR11-2, considered to be best heat tolerant, had the lowest number of its DEGs in both anthers and stipules, indicating that PR11-2 might be able to maintain a relatively steady transcriptome when subjected to short term HS. In anthers, PR11-90 had a similar number of total DEGs as that of CDC Amarillo, but CDC Amarillo had a greater number of up-regulated genes and a less number of down-regulated genes than PR11-90. Whereas in stipules, CDC Amarillo had both higher number of up-regulated and down-regulated genes than PR11-90. It is worth noting that CDC Amarillo appeared to have more unique DEGs in heat response compared with the other two varieties whose genetic backgrounds were more similar. This finding implied that heat response could depend on genetic variability.

### GO grouping on common DEGs among varieties

With the purpose of characterizing a general pea plant heat response, GO enrichment analysis was conducted on the common DEGs among the three pea varieties in this study. In anthers, GO terms relating to the 588 common up-regulated genes and 220 down-regulated genes were tested separately against the pea reference transcriptome (Pisum_sativum_v1a_GO, database was retrieved in November, 2020) to identify the significantly over-represented GO terms in biological processes under HS. All significant GO terms were filtered via hypergeometric test method at FDR adjusted p value≤0.01. Respective analysis was similarly conducted on the common genes in stipules as well, i.e., 463 DEGs with upregulation and 416 DEGs with downregulation. Up-regulated genes were enriched with 31 and 13 GO terms in biological processes for anthers and stipules, respectively (Fig 4, S1 Table). The top 10 most significant GO terms in anthers were protein folding (GO:0006457, 21 enriched terms), embryo development (GO:0009790, 9), multicellular organismal process (GO:0032501, 17), response to heat (GO:0009408, 5), multicellular organism development (GO:0007275, 15), galactose metabolic process (GO:0006012, 4), regulation of transcription, DNA-templated (GO:0045449, 43), regulation of cellular metabolic process (GO:0031323, 44), regulation of RNA metabolic process (GO:0051252, 26), and regulation of gene expression (GO:0010468, 44). In the stipules located on the same anther bearing node, the ten most over-represented GO terms were protein folding (GO:0006457, 13), response to heat (GO:0009408, 4), cellular protein modification process (GO:0006464, 12), carbohydrate metabolic process (GO:0005975, 80), post-translational protein modification (GO:0043687, 12), transcription, DNA-templated (GO:0006351, 21), RNA biosynthetic process (GO:0032774, 19), phosphate-containing compound metabolic process (GO:0006796, 11), phosphorus metabolic process (GO:0006793, 11) and regulation of RNA metabolic process (GO:0051252, 18). Four GO terms were common between anthers and stipules, which were GO:0006457 (protein folding), GO:0009408 (response to heat), GO:0006351 (transcription, DNA-templated) and GO:0051252 (regulation of RNA metabolic process). GO:0051252 is one of the ancestor terms of GO:0006351 in the cluster. Other enriched GO terms of consistently up-regulated genes in anthers were involved with primary metabolic processes, cellular respiration and reproductive structure development and the regulations of several biosynthetic and metabolic clusters including cellular metabolic and biosynthetic process, RNA metabolic process, macromolecule metabolic and biosynthetic process (S1 Table).

**Fig 4.**
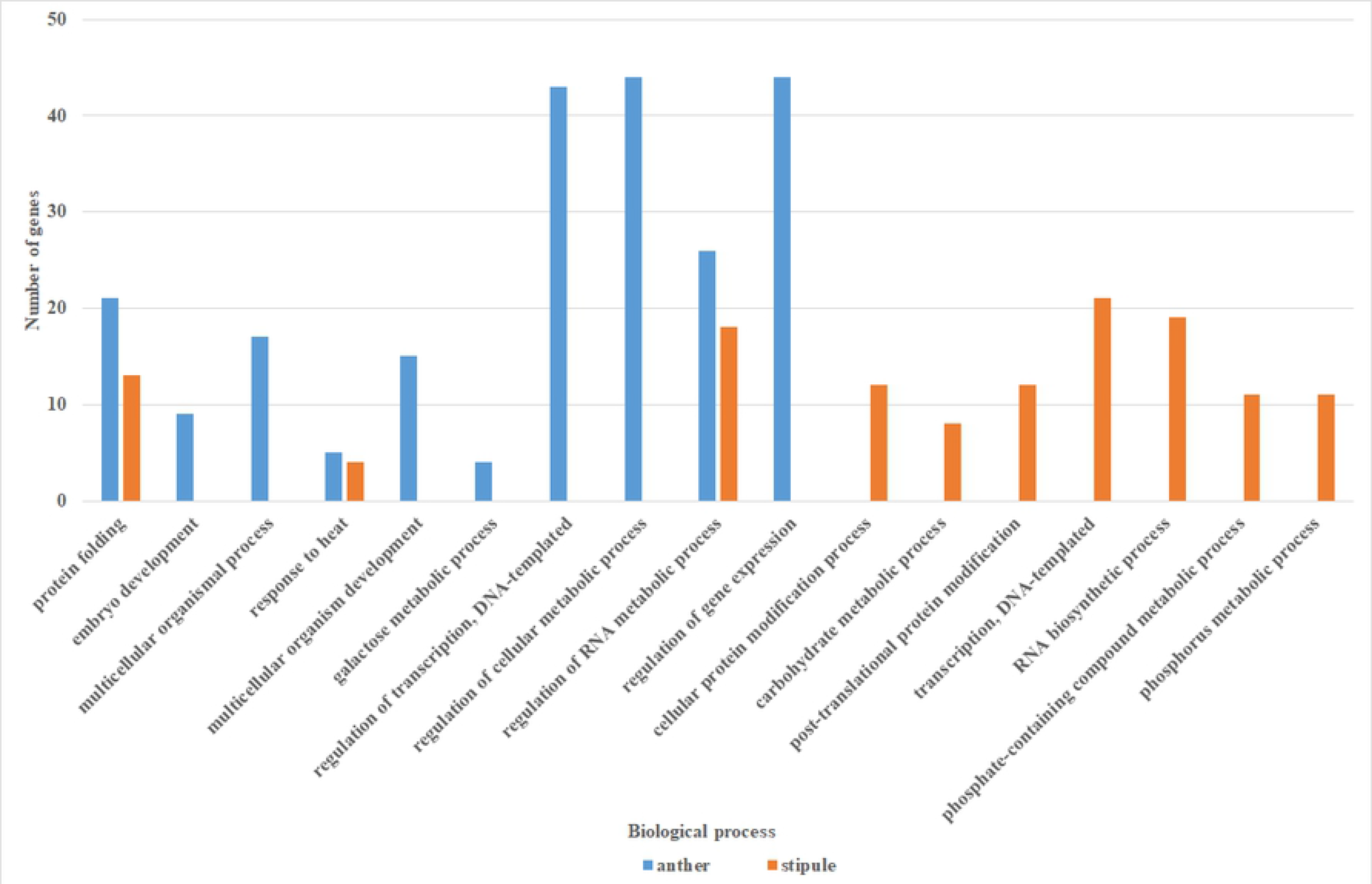
Top ten over-representative GO terms with up-regulation at log2 FC >2 in biological process in anthers (blue column) and stipules (orange column).

To further compare the most heat responsive genes among different varieties, we arbitrarily filtered the DEGs of each variety within the top 20% fold threshold range and found the most heat inducible genes were quite similar, though the greatest gene expression fold threshold varied slightly among the three varieties. The gene group relating to heat shock transcription factor (HSF) and heat shock protein (HSP) accounted for a large proportion.

Many of these HSF and HSP genes were reported here for the first time; HSF genes in particular, which expanded the previously limited findings. Putative HSF family A and B genes appeared to heat inducible, these genes included three HSF A genes and two HSF B genes (corresponding gene locus refers to Table 4). In addition to the two pea HSP genes that were previously documented, 11 other small HSP genes on chromosomes II, IV, V, VI and VII were highly heat inducible among all varieties regardless of plant organs. Increased expression of six HSP70 genes, two HSP90 family genes and three other HSP genes were also identified among all three varieties. Several heat shock cognate genes (HSCs), whose expression was previously considered as constitutive during normal plant development, appeared to be heat responsive as well (e.g. PsHSC 71.0, HSC 70-2 like etc). The response of several other genes whose functions closely interacted with HSP were also detected in this study.

**Table 4.** List of pea HSP and HSF related genes that were induced in response to 3 h 38°C heat treatment; numbers in the table are averaged log2 FC across three biological replicates.

| Locus ID | PR11-2 |  | PR11-90 |  | CDC Amarillo |  | Gene annotation |
| --- | --- | --- | --- | --- | --- | --- | --- |
|  | anther | stipule | anther | stipule | anther | stipule |  |
| Psat0s1635g0080 | 8.8 | 8.1 | 9.0 | 8.7 | 8.2 | 8.0 | PsHSP18.1 |
| Psat2g036480 | 10.8 | 10.4 | 12.5 | 9.3 | 11.4 | 9.1 | PsHSP17.9 fragment |
| Psat0s3930g0040 | 10.9 |  | 12.1 |  | 11.7 | 8.2 | PsHSP71.2 |
| Psat3g049640 | 2.8 |  | 2.9 | 2.3 | 2.9 | 2.3 | PsHSC71.0 |
| Psat0s3914g0040 |  |  | 2.5 |  | 2.4 |  | putative HSF |
| Psat1g102600 | 2.0 | 3.9 | 2.1 | 4.0 |  | 5.4 | HSF B-2A-like |
| Psat2g021040 | inf |  | inf | 2.4 |  |  | putative HSF |
| Psat3g061600 | 9.3 | 7.8 | 9.7 | 7.8 |  | 7.5 | HSF A3 |
| Psat4g086800 | inf |  | inf |  | inf |  | HSF24-like |
| Psat5g036400 | 7.7 | 7.4 | 9.2 | 8.1 | 7.1 | 7.5 | HSF A fragment |
| Psat6g059040 | 10.1 | 6.1 | 9.8 | 6.8 | 8.7 | 6.2 | putative HSF A3 |
| Psat6g078240 |  | 5.7 |  | 5.8 |  | 6.9 | HSF A3-like protein |
| Psat6g200480 | 5.8 | 7.1 | 7.2 | 7.9 | 5.6 | 8.1 | HSF B2A-like isoform X2 |
| Psat6g204120 |  |  |  | 3.2 |  | 2.5 | putative HSF A3 |
| Psat7g004560 |  | 2.0 |  |  |  | 2.7 | putative HSF |
| Psat7g131680 |  |  |  |  |  | 2.4 | putative HSF |
| Psat7g170680 |  |  |  |  |  | 3.6 | HSF A1B-like isoform X2 |
| Psat0s529g0040 | 10.1 | 10.0 | 12.5 | 9.0 | 12.6 | 9.0 | putative class I HSP20 |
| Psat2g046440 | 6.3 | 6.6 | 8.2 | 8.2 | 6.2 | 7.8 | HSP15.7 peroxisomal-like |
| Psat4g136720 | 7.3 | 7.4 | 8.1 | 8.5 | 7.0 | 7.4 | small HSP |
| Psat5g035320 | 10.5 | 10.3 | 11.9 | 9.3 | 12.2 | 9.3 | class II HSP17.1 |
| Psat4g166400 | 10.3 | 9.9 | 12.3 | 8.9 | 13.0 | 9.5 | cytosolic class II HSP |
| Psat5g073280 | 11.7 | 10.5 | 13.1 | 9.8 | 11.1 | 8.9 | small HSP |
| Psat5g174800 | 9.1 | 7.4 | 11.0 | 6.9 | 8.5 | 7.1 | putative HSP20 |
| Psat6g112800 | 10.8 | 10.0 | 13.0 | 9.4 | 11.8 | 9.0 | class IV HSP22.7 |
| Psat7g114760 | 7.5 | 9.9 | 11.3 | 9.0 | 11.5 | 11.9 | class I HSP17.6 |
| Psat7g115480 | 7.1 | 9.0 | 7.6 | 9.3 | 6.7 | 8.4 | HSP18.1 |
| Psat7g211720 | 5.4 | 6.4 | 6.0 | 6.9 | 4.5 | 6.2 | class I HSP17.5 |
| Psat7g255520 |  | 2.8 |  | 3.2 |  | 3.6 | HSP26.5 |
| Psat6g238960 |  |  |  |  | 2.3 |  | DnaJ/HSP40 cysteine-rich domain protein |
| Psat1g212880 | 7.1 | 4.9 | 8.1 | 5.7 | 7.1 | 6.5 | HSP70, mitochondrial |
| Psat1g222760 | 4.9 |  | 5.2 |  | 4.5 |  | Stromal HSP70 |
| Psat2g051360 | 8.9 | 6.5 | 10.4 | 6.6 | 8.4 | 7.2 | HSP70 |
| Psat3g143400 |  | 2.5 | 2.1 | 2.6 |  | 3.5 | HSP70-interacting protein |
| Psat3g180040 |  | 2.0 |  | 2.4 |  | 2.2 | HSC70 2-like |
| Psat3g183720 | 5.1 | 5.9 | 5.4 | 6.5 | 4.6 | 6.1 | putative HSP70 family |
| Psat4g003160 |  |  | 2.1 |  |  |  | HSP70-interacting protein |
| Psat4g035840 |  |  |  | 2.2 |  |  | HSP70 8-like |
| Psat4g210520 | inf | 3.9 | 6.5 | 4.2 | 5.4 | 5.0 | HSP70 |
| Psat5g299000 | 3.8 | 2.9 | 4.3 | 3.1 | 3.6 | 3.8 | putative HSP70 family |
| Psat7g023360 | 4.5 | 4.3 | 5.2 | 4.9 | 4.5 | 4.8 | HSP70 |
| Psat7g218840 | 8.9 | 7.7 | 10.0 | 8.3 | 8.6 | 7.8 | HSC70 2-like |
| Psat7g237280 | 4.7 | 4.4 | 5.1 | 4.9 | 4.7 | 5.3 | HSP70 nucleotide exchange factor FES1-like |
| Psat2g006440 | 5.4 | 4.6 | 5.7 | 5.4 | 5.2 | 5.4 | HSP81-2 |
| Psat0ss29864g0040 | 10.8 | 10.2 | 12.1 | 9.1 | 13.0 | 9.2 | HSP83-like fragment |
| Psat3g104360 | 8.5 |  | 9.9 |  | 8.3 |  | HSP83-like fragment |
| Psat5g164840 | 2.6 | 2.0 | 3.0 | 2.8 | 2.5 | 2.8 | HSP80 cognate protein |
| Psat6g123080 | 4.2 | 3.8 | 4.7 | 4.2 | 3.9 | 4.9 | activator of HSP90 ATPase homolog 1-like |
| Psat2g178800 | 5.0 | 3.8 | 5.5 | 4.0 | 4.9 | 4.6 | HSP70-HSP90 organizing protein 3-like |
| Psat3g067000 | 5.2 | 3.9 | 5.7 | 4.4 | 4.9 | 5.2 | activator of HSP90 ATPase homolog 1 |
| Psat0s3618g0080 | 2.4 | 3.4 | 3.0 | 4.0 | 2.2 | 5.2 | HSP DnaJ; putative transcription factor C2H2 family |
| Psat1g204360 | 10.3 | 10.2 | 11.7 | 9.2 | 12.5 | 9.4 | HSP DnaJ |
| Psat2g037160 | 3.3 |  |  |  |  |  | HSP |
| Psat5g035280 | 10.7 | 10.4 | 12.0 | 9.4 | 10.9 | 9.2 | Class II HSP |
| Psat5g229840 | 3.8 |  | 3.7 |  | 2.3 |  | Class IV HSP |
| Psat6g021840 | 9.5 | 8.2 | 11.1 | 8.8 | 10.4 | 7.5 | Class II HSP |
| Psat7g114360 | 10.6 | 10.2 | 12.5 | 9.4 | 10.9 | 9.0 | HSP |

Note: for the cells denoting ‘inf’ as FC, the reason was that their transcript at control temperature was too low to quantify. Because their transcript at HS were significant, they were still considered as heat responsive.

Interestingly, several HSP genes only responded in one organ. Anthers had unique HSFs (Psat0s3914g0040, putative HSF; Psat4g086800, HSF24-like), three HSP genes which were Psat1g222760 (Stromal HSP70), Psat3g104360 (HSP83-like fragment), Psat5g229840 (class IV HSP). Stipules had unique HSF (Psat6g078240, HSF A3-like) and two HSP genes (Psat7g255520, HSP26.5; Psat3g180040, HSC70 2-like). Several other HSF and HSP genes were specific to variety, e.g., two HSP relating genes (Psat4g003160 and Psat4g035840) were only induced in PR11-90.

A total of 220 commonly down-regulated genes in anthers among the three varieties were enriched in 18 GO terms in biological process category and 416 consistently down-regulated genes in stipules had 16 GO terms significantly over-represented (Fig 5). Ten GO terms overlapped between the two organ types, that is, GO:0006629 (lipid metabolic process), GO:0006869 (lipid transport), GO:0010876 (lipid localization), GO:0044036 (cell wall macromolecule metabolic process), GO:0071554 (cell wall organization or biogenesis), GO:0006979 (response to oxidative stress), GO:0005975 (carbohydrate metabolic process), GO:0006022 (aminoglycan metabolic process), GO:0043086 (negative regulation of catalytic activity), and GO:0044092 (negative regulation of molecular function). GO:0006508 (proteolysis), GO:0006468 (protein phosphorylation), GO:0015833 (peptide transport) and GO:0006857 (oligopeptide transport) were distinctly enriched in stipule down-regulated genes, whereas GO:0005976 (polysaccharide metabolic process), GO:0010383 (cell wall polysaccharide metabolic process), GO:0042545 (cell wall medication) and GO:0071555 (cell wall organization) were only enriched in anthers. Although more than half of the over-represented GO terms overlapped between heat stressed pea anthers and stipules at the same flowering node, surprisingly, the gene composition relating to these biological processes varied between the two organs. For example, three GO terms related to lipid biological processes were both down-regulated in anthers and stipules. However, only two genes (*PsLTP1* and *PsLTP2*) for lipid transport/localization were common, and seven genes (Psat1g060840, Psat1g082320, Psat1g085080, Psat2g027880, Psat3g005680, Psat5g104040, and Psat5g295040) for lipid metabolic processes were common between the two organ types (Table 5).

**Table 5.**
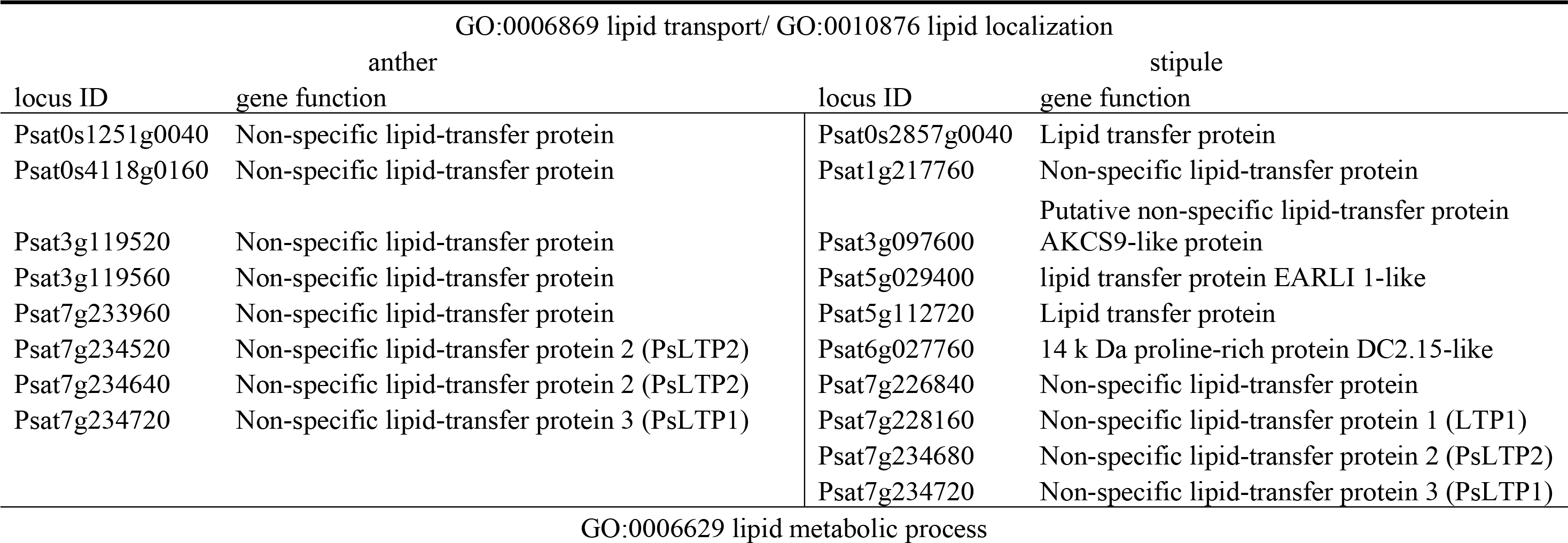
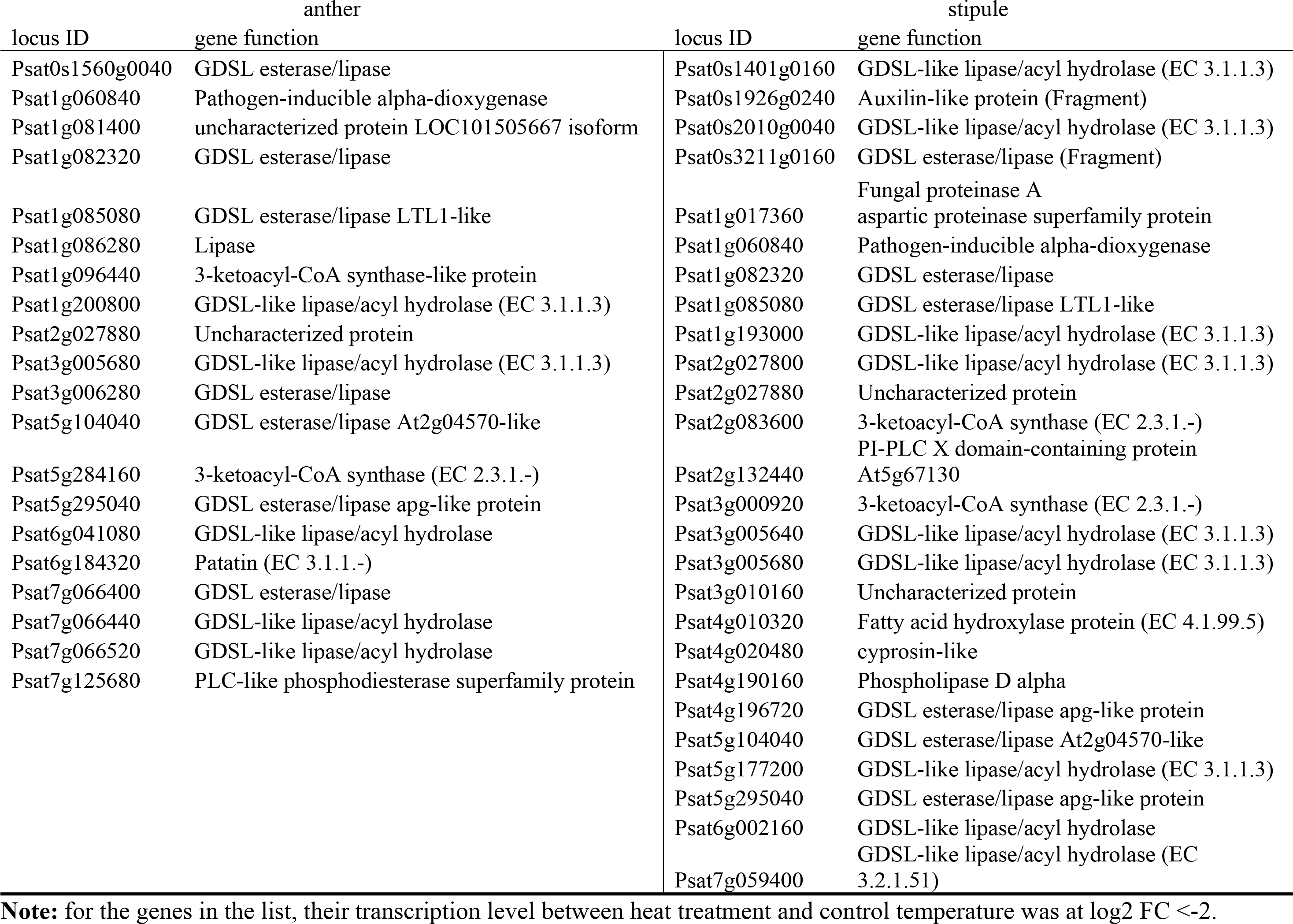
Gene locus and function list of commonly down-regulated genes that are associated with lipid transport, localization and metabolic process among the three pea varieties in anthers and stipules.

**Fig 5.**
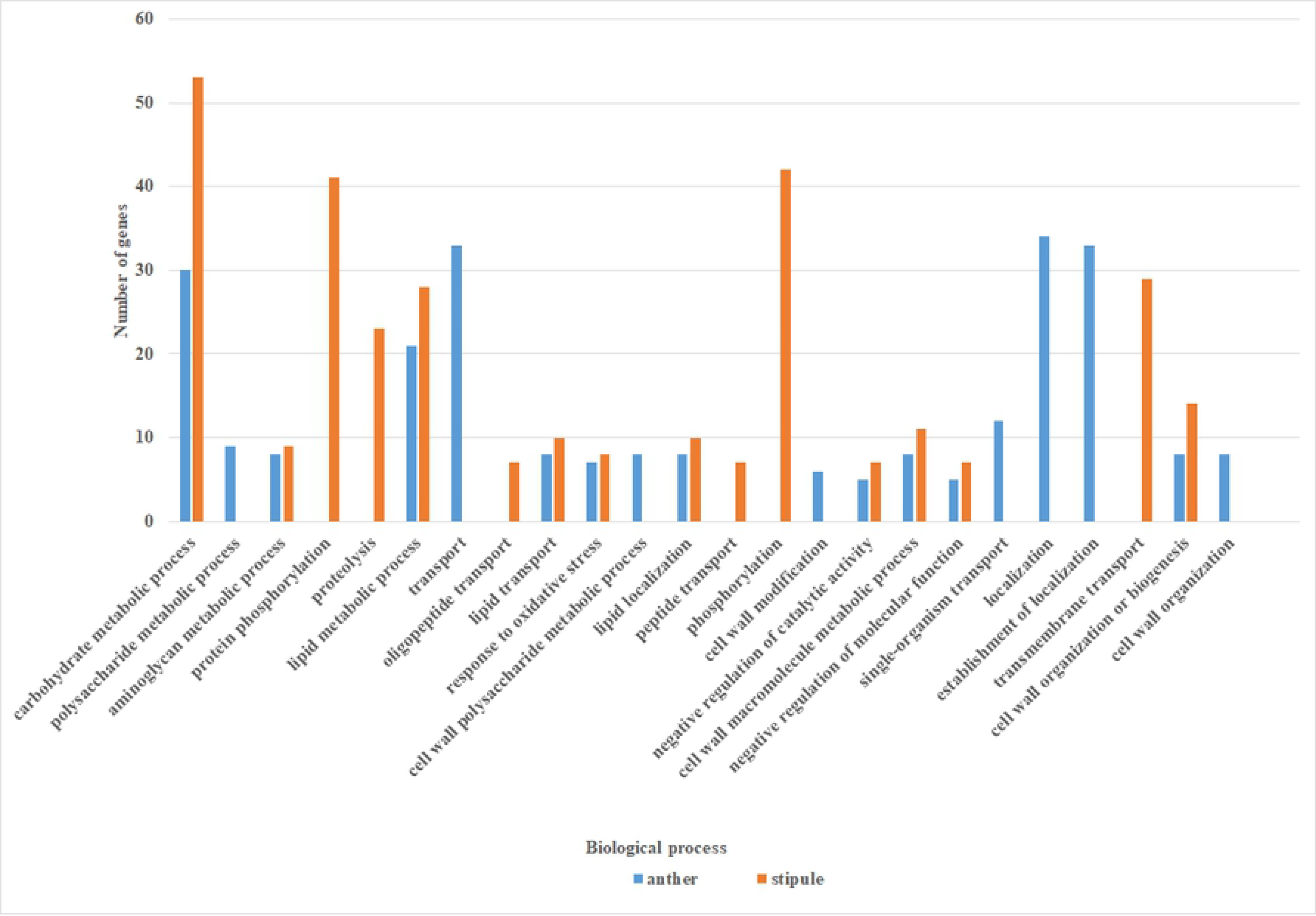
Significant GO terms (FDR adjusted p value at 0.01) in biological process of down-regulated genes at log2 FC <-2 in anther (blue column) and stipule (orange column)

We arbitrarily filtered the DEGs of each variety within the top 20% fold threshold range to further characterize a list of genes whose functions were most inhibited in heat stress. In anthers, 35 genes were shared among the three varieties, among which seven genes were involved in pectin metabolism, and three with lipid metabolism. Pectin, a polysaccharide polymer of galacturonic acid with different degrees of esterification via an a-1, 4-glycosidic bond, is a primary composition in the plant cell wall and cell interlayer. In stipules, 51 genes were common among the three varieties. The functions of these genes seemed various, including four lipase genes. Only one gene at locus Psat3g196040 was present in both lists.

### GO analysis on variety-dependant DEGs

To compare heat response among the three pea varieties, individual GO enrichment analyses were performed on the distinct DEGs of each variety, which were exclusive DEGs from the varieties’ common DEGs in individual variety DEG list. Among the three varieties, PR11-2 had the lowest number of enriched GO terms in down-regulated genes and the highest number of over-representative GO terms in up-regulated genes of both anthers and stipules, implying that PR11-2 is likely to have a superior heat tolerance compared to the other two varieties (Fig 6). In the anther transcriptome of PR11-2, no GO term was significantly enriched for down-regulated DEGs but 13 terms were up-regulated. These terms corresponded to four biological pathways, i.e., cell respiratory electron transport chain (GO:0022904), cell wall lignin metabolic and catabolic process (GO:0009808, GO:0046274), oxidation-reduction process (GO:0055114), and cellular modified amino acid catabolic process (GO:0042219). In contrast, the up-regulated GO terms in its stipule were related to regulation of transcription (GO:0006355), DNA repair (GO:0006281), and response to hormone (GO:0009725).

**Fig 6.**
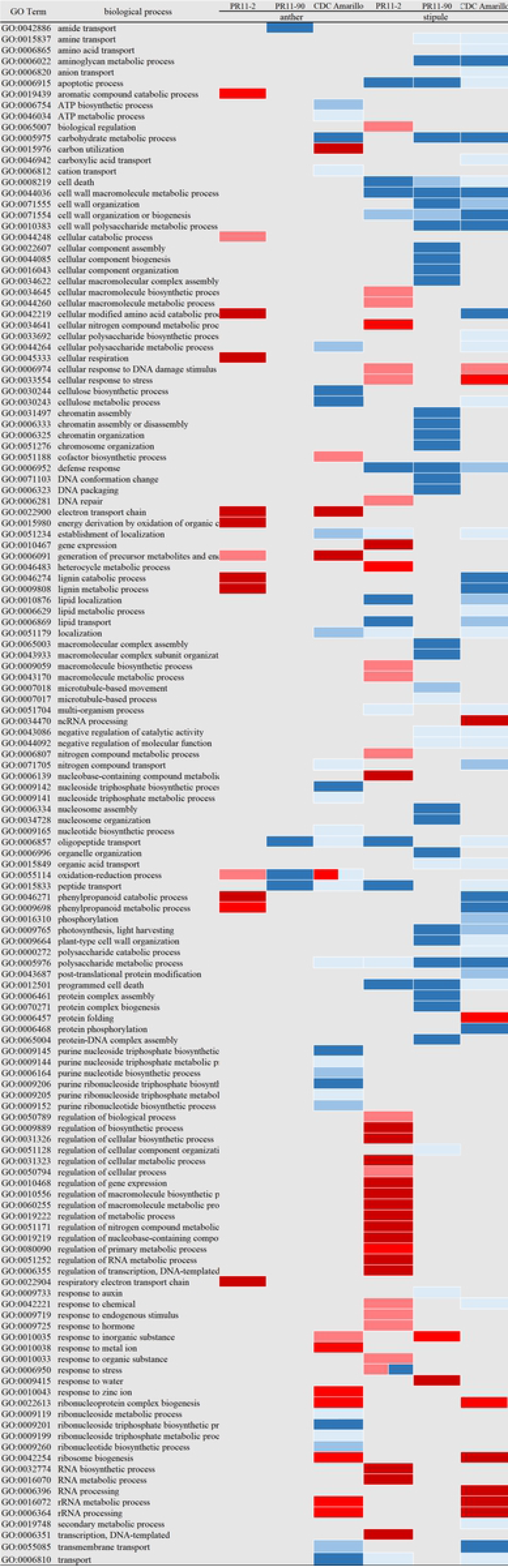
Enriched GO terms in biological processes of variety distinct DEGs in anthers and stipules. Red color are up-regulated processes, and blue color is down-regulated processes. More intense color means greater significance. Up-regulated biological process is colored in red, red scale for significance p value is shown as follows, 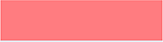 0.01≤p<0.001 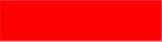 0.001≤p<0.0001 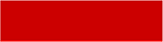 ≤0.0001. Down-regulated biological process is colored in blue, blue scale for significance p value is shown as follows, 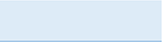 0.01≤p<0.001, 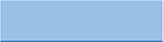 0.001≤p<0.0001, 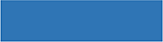 ≤0.0001.

PR11-90 (heat susceptible) had none and two GOs significantly up-regulated in anthers and stipules, respectively; whereas four and 39 GO terms were down-regulated in anther and stipule. The two up-regulated terms corresponded with response to water (GO:0009415). The four anther down-regulated terms were associated with amide transport (GO:0042886), oligopeptide transport (GO:0006857) and oxidation-reduction process. And the 39 GOs with down-regulation in stipule included 19 terms in the cluster of nucleosome assembly (GO:0006334), two terms in microtubule-based movement (GO:0007018) and response to auxin (GO:0009733) that were only over-representative in PR11-90. These distinctly heat prohibited processes in PR11-90 were predicted to link with its heat susceptible property.

In both anther and stipule transcriptomes of CDC Amarillo, the number of down-regulated GO terms was also greater than that of GO terms with up-regulation (29/12 downregulation/upregulation in anther; 46/9 downregulation/upregulation in stipule) and had the highest total number of GOs in both anthers and stipules among the three pea varieties. This differential of heat responsive GO among the three varieties demonstrated the genetic variation of field pea in heat response and shed a light in deciphering molecular mechanism involved in pea heat response and heat tolerance. In anthers, the significantly enriched GO terms in transcriptionally inhibited genes consisted of many GOs in the cluster ATP biosynthetic (GO:0006754) and metabolic (GO:0046034) process, which was uniquely observed in CDC Amarillo. The 12 enriched terms of genes, whose expression was induced, were associated with rRNA processing (GO:0006364), response to zinc ion (GO:0010043), electron transport chain (GO:0022900) and carbon utilization (GO:0015976). rRNA processing was also up-regulated in stipule in addition to cellular response to DNA damage stimulus (GO:0006974) and protein folding (GO:0006457). The stipule down-regulated GO terms were mainly linked with amino acid transport (GO:00068650), cell wall polysaccharide metabolic process (GO:0010383), lignin metabolic and catabolic process, lipid transport (GO:0006869), lipid localization (GO:0010876), lipid metabolic process (GO:0006629) and protein phosphorylation (GO:0006468).

In stipules, 7 GO terms were down-regulated in all three varieties and were involved in apoptotic process (GO:0006915), defense response (GO:0006952) cell wall macromolecule metabolic process (GO:0044036) and polysaccharide metabolic process (GO:0005976). Cell wall polysaccharide metabolic process, plant-type cell wall organization (GO:0009664), photosynthesis, light harvesting (GO:0009765), amine transport (GO:0015837) and aminoglycan metabolic process (GO:0006022) were down-regulated in PR11-90 and CDC Amarillo. Lipid transport and localization were down-regulated in PR11-2 and CDC Amarillo. It is noted that cellular response to stress (GO:0033554) and cellular response to DNA damage stimulus (GO:0006974) was only up-regulated in the stipules of two heat-tolerant varieties, PR11-2 and CDC Amarillo. Comparing the between the two varieties, transcripts of four genome loci were common, which were Psat2g148040, Psat5g135640, Psat6g105320 and Psat6g199840. In anthers, generation of precursor metabolites and energy (GO:0006091) and electron transport chain (GO:0022900) were only up-regulated in PR11-2 and CDC Amarillo as well, and the transcriptional level of three relating genes (Psat1g132320, Psat1g132440, and Psat6g041400) was induced in both varieties. Interestingly, contrasting response was observed in oxidation-reduction process (GO:0055114) among the anthers’ transcriptomes of the three varieties. This biological process was enriched in the up-regulated genes in PR11-2, but significant in the down-regulated genes in PR11-90 and were over-representative in both down/up-regulated genes in CDC Amarillo. Among these genes, 8 genes were commonly up-regulated in PR11-2 and CDC Amarillo, 28 genes were down-regulated in CDC Amarillo and PR11-90 (S2 Table).

## Discussion

### General and genotype specific heat response at cellular level

Separate heat responsive genes of individual variety were identified at log2 FC >2 for anther and stipules, and two heat tolerant varieties in our study demonstrated different transcriptomic response, i.e., PR11-2 had the lowest number of DEGs among the three varieties, contrastingly DEG number in CDC Amarillo was the greatest. This was also seen in maize, where tolerant cultivar S058 and L043 had the most and least abundant DEGs among four tolerant and four susceptible varieties, respectively [16]. Collectively, it is suggested that plant heat tolerance could be achieve in different mechanism.

Individual GO enrichment analysis was carried out on common DEGs among the three varieties and variety unique DEGs, aiming to characterize the general heat response in biological processes of pea plant as well as unique responses relating to heat tolerance. Response to heat (GO:0009408), protein folding (GO: 0006457) and transcription, DNA-templated (GO:0006351 and GO:0051252) were commonly upregulated between stipules and anthers (Fig 4). The transcriptome re-program and chaperone function of HSPs are considered to contribute to plant’s basal thermo-tolerance [41]. Regressed biological processes were mainly related to lipid transport, lipid metabolic process and cell wall macromolecule metabolic process, and their relevance to HS and relating genes are further discussed in latter section.

Regarding variety unique heat response, anther of PR11-2 had only up-regulated processes, belonging to three biological process clusters, i.e., respiratory electron transport chain, lignin catabolic process and cellular modified amino acid catabolic process (Fig 6). PR11-90 had none induced biological process in anther. This could partly explain heat tolerance of PR11-2 over PR11-90. Intriguingly, electron transport chain (ETC) was also upregulated in CDC Amarillo. In electron transport chain, Psat1g132320 and 6g041400 encoding mitochondrial cytochrome b and Psat1g132440 encoding uncharacterized protein were upregulated. Cytochrome b-c1 complex is an essential component of the mitochondrial ETC. Chilling induced accumulation of reactive oxygen species resulting from over-reduction of ETC led to oxidative stress [42].

In stipules, cellular response to DNA damage stimulus was only induced in two heat tolerant varieties. Four genes were common between gene lists of the two varieties, which were 2g148040 (DNA mismatch repair protein MLH3), 5g135640 (DNA excision repair protein), 6g105320 (cryptochrome 2b), and 6g199840 (DNA mismatch repair protein MSH3). The putative functions of the four genes were involved with three DNA repair pathways, but these pathways were well studied in UV light induced stress [43]. Elucidation on the connection of the plant DNA repair to abiotic stress responses remains scarce, plant’s ability to maintain its genome integrity is likely to play a role in stress tolerance [44].

### Regulatory importance of HSFA3 and HSFB2 in heat response

Although HSFs are believed to play a central regulation role in the transcriptional induction of downstream HS responsive genes, HSFs display their variation in HS response in terms of induction fold threshold and regulation, and thereby could affect various gene expression induction. Structurally plant HSFs are classified into three classes, namely, HSF A, B, and C, based on their structural peculiarities. The best characterized HSF gene family in plants has been firstly reported in Arabidopsis (21 HSF genes) [9]. Wheat (56 HSF genes) [45]; and soybean (52 HSF genes) [46] were reported to have the largest families in monocot and dicot crops, respectively. Among the three classes, the function of HSFAs was more clearly elucidated, and here is broad agreement that their role most directly leads to heat-induced activation of heat shock genes. HSFA1s are predicted to be the “master regulators” that have the direct role in the activation of transcriptional networks. Knockdown of HSFA1 genes in Arabidopsis led to a reduced induction of many HS-responsive genes, as a result plants demonstrated HS susceptible phenotypes [47, 48]. The thermo-tolerance conferred by Arabidopsis HSFA1d was further confirmed in a recent study in pea [49], where transformant pea plants with this Arabidopsis HSF was more heat tolerant than its wild type due to the increased antioxidant enzyme activity and reduced hydrogen per oxide. Another study in Arabidopsis concluded that HSFA3 was also an important HS-responsive TF, because knockout or knockdown mutation of HSFA3 resulted in reduced expression of putative target HSP genes during HS [50]. OsHSFA3 and A2s were identified to responsive in rice panicle when exposed to multiple hours of HS [51]. In comparison, in common wheat (*Triticum aestivum* L.), HSFA2 and A6 had the highest transcriptional induction among 56 TaHSF members when subjected to HS, which revealed the regulatory importance of these two subclasses during HS [45]. Among legume plants, over-expression of soybean *GmHSFA1* could enhance the thermotolerance of transgenic soybeans via the activation of various HSP gene expression [52]. In the other study, the induction of GmHSFs at HS was found to variate at different plant stages, including HSFA1 [53]. In *Lotus japonicus*, HSFA1 did not dominantly express in heat-stressed seedlings, A2, A3, A6, A7, B2 and B5 were exclusively heat induced and other hsf subclasses could also be involved in other abiotic stress responses [54]. Q-PCR expression analysis of chickpea HSFs under heat stress at pod development and at 15 days old seedling stage showed that CarHSFA2, A6, and B2 were constitutively up-regulated at both plant development stages indicating their importance in the regulatory network relative to HS [55]. In the present study, various transcripts of putative pea HSFs were characterized that were responsive to 3h heat treatment, among which putative HSFA stood out in its amount abundance, the A3 subclass in particular. Three HSFA transcripts (Psat3g061600, Psat5g036400 and Psat6g059040) were highlighted because their transcriptional levels were dominantly increased in both anthers and stipules in all three varieties (Table 4), suggesting they are essential transcriptional regulators in pea HS response. Further analysis on knock-out mutants of these HSF genes will validate their exact role, whether directly or not, in heat regulation. Interesting, individual HSF were identified for anthers and stipules, indicating different regulatory networks may exist between vegetative and reproductive organs.

Functions and molecular mechanism of HSFBs were less elucidated, but they were found to interact closely with HSFA in plant’s HS response. The role of HSFBs were reported either as a repressor or activator in the transcription of HSFA depending on plant species, as a result, they participated in different mechanisms in HS regulation. In *A. thaliana*, HSFB suppressed the transcriptional activities of HS-inducible HSFs, including HSFA2, A7a, at both normal temperature environment and HS condition [56]. On the contrary, the function of tomato’s HSFB1 seemed more complex, it could work either as a co-activator of some HSFs e.g., HSFA1a or as a transcription repressor of other HSFs such as HSFA1b and HSFA2([57–59]. In our result, transcription levels of two putative HSFB2 genes (Psat1g102600 and Psat6g200480) were highly heat induced along with HSFA genes independent of organ types and genotypes, implying their positive role in transcriptional regulation of field pea in HS, which was in agreement with the finding in chickpea [55]. It seemed that the role of HSFB in legume crops was similar to the coactivator characteristics of tomato HSFB.

### Transcriptional induction of various pea sHSPs and HSP70 at HS

In plant cellular defense against HS, the induction of HSP is one of the major responses. HSPs act as molecular chaperones which are proteins that facilitate folding of other functional proteins especially at the secondary and tertiary structure and prevent them from denaturation and aggregation during exposure to HS. Depending on the molecular size, HSPs are divided into five conserved classes: small HSPs (sHSPs), HSP60, HSP70, HSP90 and HSP100.

sHSPs range in size from 10 to 42 kDa and share a conserved C-terminal domain that is common to all eukaryotic organisms. Generally, sHSP functions as a molecular chaperone and protects the substrate proteins against thermal aggregation or denaturation. In six legume species, more than 5 different sHSPs were detected from plant tissues exposed to HS [60]. In pea, several sHSPs belonging to two classes based on their sequence alignment and immunological cross-reactivity were isolated. *PsHSP 17.7, 17.9, 18.1* were located in the cytoplasm, whereas *PsHSP 21 and PsHSP 22* were located in chloroplasts and mitochondria, respectively ([19, 20, 61]. From these reports, we could conclude that they were all involved in establishing cellular thermotolerance to some degree, though the induction of their expression was triggered at different temperatures.

Transcriptome profiling in our experiment revealed that the transcriptional levels of cytoplasmic sHSPs were drastically increased at HS among the three pea varieties (Table 4), which was in agreement with the above-mentioned result on other pea genotypes, suggesting the function of these sHSPs is general in field pea plant. Beyond that, transcriptional response of other sHSPs in relation to HS were also characterized, which provides a more comprehensive picture of sHSP heat response in pea.

HSP70 proteins have also been extensively studied; they are ATP-driven molecular chaperones with an N-terminal ATPase domain and a C-terminal peptide binding domain. Similar to the gene family encoding sHSPs, HSP70 genes also encode proteins targeted to different cellular compartments, including mitochondria, chloroplast, endoplasmic reticulum, and the cytoplasm. Similarly, HSPs isolated in pea differed in their expression under different temperature environments, inferring functional differences between heat-induced and constitutively expressed HSP 70 homologues. In our study we confirmed the significance of various HSP70 genes in field pea heat response.

### HS response in pea cell wall

Various biological processes relating to cell wall were significantly down-regulated when exposed to HS in our study, which helped decipher the molecular mechanism of heat damage on pea cell wall (Fig 5 & 6). Similar in heat stressed lentil, a major group of heat responsive genes were involved in plasma membrane and cell wall [18].

Plant cell walls have multiple layers and are made up of three sections, i.e., the middle lamella, primary cell wall, and secondary cell wall. The primary wall surrounds growing cells or cells capable of cell growth and its plasticity is essential for cell expansion and growth; whereas the secondary wall is a highly specialized and thickened structure to provide the sufficient rigidity, which undergoes irreversible changes in many fully developed cells. The middle lamella is a pectin layer to provide necessary adhesive between two adjoining cells [62]. Pectin, a mixture of polysaccharides, is also a major composition in primary cell wall, especially in dicotyledonous plants [63]. In addition to its adhesive property, adjustment of its content in cell wall is proposed to link with various physiological function during plant life cycle as well as contribute to signal transduction to various conditions. Reproductive tissues are particularly rich in pectin compared with other tissues, for example pectin constituted ∼40% and 15% in rice pistil and anther cell wall, respectively, whereas the proportion of pectin was only 5% in the cell wall of mature leaf [64]. Transcriptome comparison of this study between HS and normal temperature characterized a cluster of genes encoding pectin esterase (enzymes for pectin metabolism), only heat responsive in anthers of all three varieties, not in stipule, and it is proposed to be associated with contrasting pectin composition between reproductive organ and vegetative plant organ. The reduced expression of pectin methyl esterase (PME; EC 3.1.1.11) genes under HS was consistent with the finding in canola [17]. Intriguingly, recent studies in pea aluminum stress and cold stress suggested that the degree of pectin methyl-esterification and PME activity could also play a role in both abiotic stresses [65, 66]. Still, the stress effect on the architecture of cell wall remodeling by PME activity may depend on the plant species, genotype, and growth stage, and also rely on the intensity and timing of the stress [62].

Lignin is a major composition in secondary cell wall and provides cell structural rigidity. Its biosynthesis consists of a very complicated network, where cinnamyl alcohol dehydrogenase (CAD), laccase (LAC) and peroxidase are involved. In *A.thaliana*, CAD function defective mutant displayed inhibited plant and male sterile compared with wild type, likely attributed to the abnormally reduced lignin biosynthesis in the anther [67]. Likewise, CAD1 mutant of *M. truncatula* had a much lower lignin content than the wild type, though causing no growth difference between two materials at normal temperature environment (22°C), the growth of this *MtCAD1* mutant was significantly suppressed at 30°C [68]. In our study, lignin metabolic and catabolic process was identified to be uniquely up-regulated in the anther’s transcriptome of heat tolerant variety, PR11-2, when exposed to HS (Fig 6). The genes in this process were identified to be LAC encoding genes on pea chromosome II, III, V and VII, which are predicted to be associated with heat tolerance. In Anadiplosis, functions of LAC 1, 4 and 17 were linked with anther dehiscence success [69]. A QTL was identified for HS susceptibility index of percent spikelet sterility in rice on chromosome XII, and one LAC gene was included in this QTL interval [70].

### Effects of HS on Lipid Transport and Metabolism

HS in our study adversely affected lipid transport and localization in both pea anther and stipule independent of genotypes (Fig 5). The lipid process was inhibited mainly via the down-regulation of various transcripts encoding non-specific lipid transfer proteins (LTPs; Table 5). Plant LTPs are broadly categorized into LTP1 and LTP2 groups based on the molecular weighs. LTP1s generally consist of 90 amino acids, whereas LTP2s have around 70 amino acids.

Although the biological functions of LTP have not been clear yet, previous studies suggested that LTPs genes can be divided into three groups depending on expression patterns of the related genes, that is, 1) genes only expressed in aerial plant parts; 2) genes only expressed in root; and 3) genes whose expression was restricted in reproductive tissues [71]. Our results added another piece of evidence to support tissue-specific expression of LTP genes, because different transcripts of LTP genes were characterized between field pea anther and stipule at normal development as well as at HS condition. Except that the two genes encoding *PsLTP1* & *2*, previously isolated in pea seeds [72], were heat responsive in both plant samples, other corresponding genes variated. To the authors’ knowledge, our work is the first to report the link between LTP genes with pea normal plant development and heat response, and their biological functions are worth being validated via mutation experiment. In wheat, *LTP3* accumulation was detected in cell membrane after HS at plant seedling and grain-filling stages, what’s more, in transgenic Arabidopsis seedling with the overexpression of *TaLTP3* was better tolerant to HS than control plants, possibly because of a less membrane injury [73].

In addition, the lipid metabolic process was negatively damaged by HS in both anther and stipule among all three pea varieties (Fig 5), which was also seen in rice heat stressed anther [15]. The damage was mostly due to that the transcriptional activity of multiple genes associated with GDSL lipase were adversely affected, although GDSL gene family was differentially expressed between anther and stipule (Table 5). Studies in this aspect are scarce in legume including pea. In the model plant *A. thaliana*, GDSL lipase gene has a family of 108 gene members, which are distributed across plant genome [74, 75]. Among them, 20 members were expressed in all tissues, and the other 16 and five members were exclusively expressed in flower and root, respectively. Mayfield et al. (2001) reported one GDSL lipase to be involved in the formation of pollen coat [76]. With the advance in omics technology, the integration of lipidome and transcriptome provides a new perspective of studying HS as shown in [77]Higashi et al. (2015).

### Coincidence of Heat Responsive Genes among Field Pea Studies

Attempts in genomic understanding of pea HS and selecting for heat tolerant varieties have started since last decade ago, benefiting from the rapid advancement in sequencing technology. However, results from individual research can not all be compared because the types of genetic markers applied were various. Our characterized heat responsive genes can be compared with a recent association mapping study[8] by Tafesse et al. (2020), as pea genome locus markers were used in their work. Twelve DEGs in our study coincided with putative candidate genes for heat responsive trait characterized in the field condition from their work (S3 Table). The response of these 12 genes fell into three patterns: 1) responsive in all tissue types among the three varieties (e.g. Psat5g303760 encoding uncharacterized protein); 2) specifically responsive to tissue type (e.g. Psat2g144160 encoding pectin acetylesterase); 3) only responsive in certain genotype (e.g. Psat2g166520 encoding putative rapid alkalinization factor). Further functional annotation of individual gene would benefit to explicit its role in HS response.

## Conclusions

Our research profiles a global transcriptome response to short term HS among different field pea varieties. Common effects of HS in biological processes are shared between anthers (reproductive organ) and stipules on the same flowering node (vegetative organ), though the involved genes in certain processes differed between the two organs (e.g., lipid transport and metabolic process). Distinct heat responses were characterized on individual pea varieties, which provides insight into molecular mechanisms of heat-tolerance response. This research supports the utilization of RNA-Seq for the identification of heat responsive genes, provides preliminary result for marker assisted selection, and is proposed to be applicable in other abiotic stress studies of pea.

## Acknowledgements

We thank Dr. Arthur Davis at Department of Biology, University of Saskatchewan, for his advice in the experimental design.

## Supporting information

S1 Table. Over-representative GO terms with significance p value in the anther’s consistently up-regulated genes among three pea varieties.

**S2 Table. Genes in GO:0055114, oxidation-reduction process was upregulated in PR11-2 and CDC Amarillo or down-regulated in PR11-90 and CDC Amarillo.**

**S3 Table. Overlapping of heat responsive genes between our study and Tafesse et al. (2020).**

Note: Trait names, SPAD: Soil plant analysis development meter for the estimation of leaf chlorophyll concentration, CT: canopy temperature, RSL: reproductive, PN: pod number. Red cell represents up-regulated gene expression at HS in our study, whereas blue cell represents a down-regulation.

S1 Fig. qPCR primer efficiency standard curves.

